# Immunopipe: A comprehensive and flexible scRNA-seq and scTCR-seq data analysis pipeline

**DOI:** 10.1101/2024.05.14.594248

**Authors:** Panwen Wang, Yue Yu, Haidong Dong, Shuwen Zhang, Zhifu Sun, Hu Zeng, Patrizia Mondello, Jean-Pierre A. Kocher, Junwen Wang, Yan W. Asmann, Yi Lin, Ying Li

## Abstract

Single-cell sequencing technologies provide us with information at the level of individual cells. The combination of single-cell RNA-seq and single-cell TCR-seq profiling enables the exploration of cell heterogeneity and T-cell receptor repertoires simultaneously. The integration of both types of data can play a crucial role in enhancing our understanding of T-cell-mediated immunity and, in turn, facilitate the advancement of immunotherapy. Here, we present immunopipe, a comprehensive and flexible pipeline to perform integrated analysis of scRNA-seq and scTCR-seq data. In addition to the command line tool, we provide a user-friendly web interface for pipeline configuration and execution monitoring, benefiting researchers without extensive programming experience. With its comprehensive functionality and ease of use, immunopipe empowers researchers to uncover valuable insights from scRNA-seq and scTCR-seq data, ultimately advancing the understanding of immune responses and immunotherapy development.

## Introduction

T cells play an essential role in the adaptive immune system and are critical for the success of immunotherapy. The immune system safeguards our bodies from a range of illnesses, including cancer, by generating a wide variety of T-cell receptors (TCRs) through V(D)J recombination. These TCRs enable T cells to recognize and target specific antigens [1]. To study the diversity and clonality of TCR repertoire at the single-cell level, single-cell TCR-sequencing (scTCR-seq) has emerged as a powerful technique [2-4]. Additionally, single-cell RNA sequencing (scRNA-seq) allows for the analysis of gene expression in individual cells, providing insights into cellular heterogeneity and functional states [5-9]. By integrating scRNA-seq and scTCR-seq data, researchers can gain a comprehensive understanding of T-cell-mediated immunity and its potential applications in immunotherapy [10-19].

Analyzing scRNA-seq and scTCR-seq data can be complicated due to the diverse nature of the data. These types of data often include substantial amounts of high-dimensional information, which requires advanced computational methods for processing and analysis [20]. Additionally, the integration of diverse types of data, such as gene expression and T-cell receptor information, adds another layer of complexity [19]. Furthermore, analyzing immune cell populations involves identifying and characterizing various cell types, which can be challenging due to the heterogeneity and plasticity of these cells [19]. Moreover, downstream analysis after integration of scRNA-seq and scTCR-seq data is strenuous since the data encompasses gene expression profiles, TCR repertoire, and clinical phenotypes, and requires careful consideration in arranging the results to ensure clarity and coherence. Therefore, meticulous planning and organization are crucial to present the results in a manner that facilitates comprehension and enhances the overall impact of the analysis.

In this work, we have developed a flexible pipeline named immunopipe that facilitates the analysis of scRNA-seq and scTCR-seq data, empowering researchers to unlock valuable insights and accelerate the development of immunotherapeutic strategies. With its user-friendly web interface and comprehensive functionality, immunopipe is accessible to researchers with varying levels of programming experience.

Immunopipe is designed as a one-stop solution for scRNA-seq and scTCR-seq data analysis starting with read count and clonotype information. It offers a range of analyses for quality control, clustering, cell type annotation, TCR repertoire analysis, and integrative analysis of both types of data. Advanced models have been developed to better understand the relationships between TCR sequence and gene expression profiles [21-23]. However, these studies primarily focus on the development of integration models, which can be used as modules in immunopipe. Alongside other analyses, immunopipe provides a more comprehensive view of the entire dataset. Some packages can perform thorough analysis for scRNA-seq data [24, 25] or scTCR-seq data [26-29]. It’s also possible to perform integrative analysis with these tools. For example, Seurat [25] can perform scRNA-seq data analysis and then integrate scTCR-seq data with the Seurat object. However, the analysis requires granular control of each step, which can be challenging for researchers without extensive programming experience. Furthermore, extra effort may also be necessary to analyze the integrated data along with clinical phenotypes. Immunopipe streamlines these analyses to offer a comprehensive solution and provides a user-friendly interface for researchers to perform the analysis with ease.

## Results

### Abstraction of the analysis for scRNA-seq and scTCR-seq data

The abstraction of the analysis enhances the robustness and flexibility of the pipeline. In general, as shown in Figure 1 (also Figure S1 and Figure S2), the common practices for analyzing scRNA-seq and scTCR-seq data are included in the pipeline. T cell selection is performed to avoid bias introduced by non-T cells. The selected T cells are then re-clustered and integrated with the clonal information. Various downstream analyses, including differential gene expression and gene set enrichment analysis, are performed for the integrated data. In addition, the pipeline includes innovative approaches from recent methodological advancements in literature.

**Figure 1.**
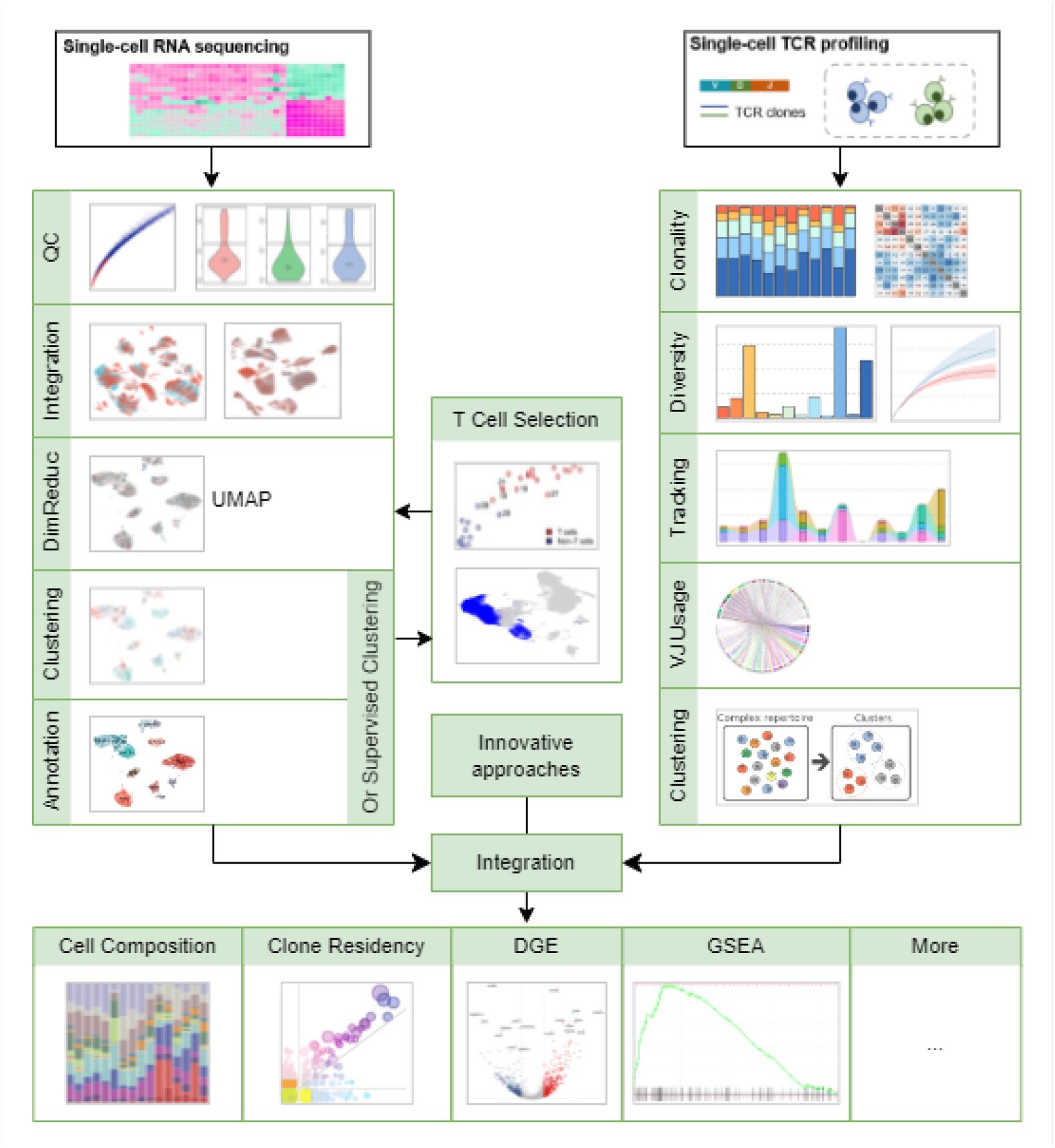
Illustration of analysis by immunopipe

### Common practices for scRNA-seq data analysis

The scRNA-seq data analysis starts with quality control to assess the overall data quality and identify potential outliers or technical artifacts. A few metrics commonly used by the community are available [30, 31], including the number of cells with expressions for a gene, and for a cell, the number of features [32, 33] and the proportion of mitochondrial, ribosomal, hemoglobin and platelet genes [30, 34]. Following this, normalization, sample integration, dimension reduction, and clustering are performed to group cells with similar expression profiles, facilitating the identification of distinct cell types or populations. The clustering process is followed by the identification of markers for each cluster and enrichment analysis using the significant markers based on their gene expression profiles.

The markers themselves and the enriched pathways can provide valuable insights into the functional states and potential roles of different cell subsets, thus facilitating cell population identification. The visualization of gene expression patterns allows researchers to explore the expression of specific genes within cell populations. Furthermore, several packages are employed to automate the cell type annotation procedure. Alternatively, cell type annotation can also be achieved using supervised clustering by mapping the cells to well-annotated reference datasets.

### Common practices for scTCR-seq data analysis

The scTCR-seq data analysis starts with basic analysis and statistics of TCR clones, such as the number of unique TCRs, clonality, and diversity metrics. Advanced analysis, such as gene usage, clonotype change tracking, expanded or enriched TCR clone identification, and repertoire overlap detection between different samples or conditions, can also be performed subsequently [35]. The clone residency analysis is useful to examine the presence and persistence of T-cell clones across different samples or time points [17]. V-J usage plots investigate the V-J gene conjunction patterns within the TCR sequences, shedding light on the repertoire diversity and potential antigen specificity [28]. Moreover, clustering based on the TCR sequences allows for the identification of T-cell clusters or refinement of clonotypes, aiding in the characterization of specific T-cell populations [36, 37].

### T cell selection

To integrate scTCR-seq data with scRNA-seq data and avoid the bias introduced by non-T cells, we provide a T-cell selection process that allows for the seamless integration of T-cell populations with scRNA-seq data. All cells are first clustered and T cells are selected based on the clonotype percentage in each cluster from scTCR-seq data and the expression of marker genes, including positive markers like CD3E, CD3D, and CD3G [38], and negative markers, or exclusive markers for other cell types, such as CD14, CD19, and CD68 [39, 40]. The selected T cells are then re-clustered and analyzed with the remaining common practices for scRNA-seq data analysis.

### Integration of scRNA-seq and scTCR-seq data

The integration of scRNA-seq and scTCR-seq data in immunopipe directly links the two types of data and derived results from them by cell IDs, so that clonal information and TCR clusters can be used in the downstream analysis, such as finding markers for specific TCR clones or TCR clusters. With the cell type annotation based on scRNA-seq data, we can investigate the TCR repertoire of different cell types. For example, we can compare the diversity of TCR clones between different cell types or identify the TCR clones that are specific to a given cell type. In addition, it is possible to perform further analysis that involves clinical phenotypes, such as immunotherapy response. This type of analysis can help identify clones that are associated with specific clinical phenotypes, which is valuable in the development of therapeutic strategies.

### Differential gene expression (DGE) analysis and gene set enrichment analysis (GSEA)

DGE analysis is a common practice in scRNA-seq data analysis. It allows for the identification of genes that are differentially expressed between two groups of cells, providing insights into the functional differences between these groups [41]. It is generally used to identify markers for specific cell types or groups. GSEA is a computational method that determines whether a priori-defined set of genes, such as pathways, shows statistically significant and concordant differences between two biological states. It is widely used to gain deeper insights into the functional states of cells and the underlying biological mechanisms [42]. Both DGE analysis and GSEA are available in immunopipe, allowing researchers to gain a more comprehensive understanding of the functional states of cells.

### Incorporation of innovative approaches

In addition to the common practices for scRNA-seq and scTCR-seq data analysis, immunopipe incorporates innovative approaches that were introduced in recent publications. Metabolic landscape analysis at single-cell resolution [43] is designed to study metabolic programs in single cells. Physiochemical attribute analysis of the CDR3 amino acid sequences can be performed by comparing two groups of cells, typically regulatory T cells and resting conventional T cells. It quantifies hydrophobicity, isoelectric point and volume of the amino acids in the CDR3, which are associated with self-reactivity of the TCR [44-47]. TESSA [23] characterizes TCR repertoire by embedding TCR sequences and integrating them with scRNA-seq data, which has shown superiority over methods using TCR sequences only, in terms of the association of TCR clones with clinical phenotypes.

### Flexible configuration and execution options

The flexibility of Immunopipe is reflected at different levels, from the overall workflow based on input data to the detailed configuration of specific analyses. Although designed to analyze both scRNA-seq and scTCR-seq data (Figure S1), immunopipe can be used for scRNA-seq data analysis alone (Figure S2). The essential analyses that adhere to common practices are mandatory; while most of the other analyses are optional. This provides users with an elevated flexibility to suit their individual needs. T cell selection, for example, is not necessary if pure T cells are sorted in scRNA-seq data. Immunopipe also offers granular control of each analysis, allowing users to adjust the parameters to suit their specific design. Many of these parameters are pre-configured with carefully selected values, thereby facilitating the analysis process adaptable and straightforward.

In addition to gene expression profiling by scRNA-seq data, the analysis performed by immunopipe heavily relies on the cell grouping information within the data that could be retrieved from scTCR-seq and clinical data. Users have the option to directly reference the relevant variable from the metadata in the configuration, aligning with their specific research requirements. Additionally, it is also possible to derive new variables in the configuration from existing ones by modifying the metadata, which can subsequently serve as grouping information for the analysis, adding another layer of flexibility. Moreover, data can be filtered through configuration, which enables focusing on a subset of cells that are of interest. It is frequently necessary to conduct similar analyses multiple times. For instance, multiple variables may need to be overlayed on a dimension reduction plot or sub-clustering is desired to perform on multiple subsets of cells. The configuration scheme is designed to accommodate such scenarios, enabling users to effectively perform these analyses and explore data from various perspectives with ease and efficiency.

The pipeline is executed on a local machine by default, while also offering the possibility of running it on alternative scheduler platforms. Immunopipe is equipped with built-in support for executing the pipeline via Sun Grid Engine, Slurm Workload Manager, and SSH. The entire pipeline, along with its dependent packages, is compiled into a docker image (see Code availability), enabling seamless deployment across diverse computing environments. This ensures flexibility and compatibility, allowing users to execute immunopipe on their preferred platforms. Furthermore, the use of a docker image simplifies the installation procedure and mitigates the potential for dependency conflicts which can be a big hurdle for less experienced users.

The results generated by immunopipe are organized and presented comprehensively and intuitively. The pipeline generates reports in HTML format, which can be easily delivered or hosted on a server for easy access and interpretation. The report is structured by modules, and the results of each module are presented in a variety of formats, including tables and plots.

Immunopipe is also accompanied by a web interface, which facilitates effortless configuration file generation, and initiation and monitoring of the execution. This interface streamlines the analysis process and provides convenient access to log files, intermediate results, and final reports. The web interface offers detailed descriptions of each configuration option, allowing users to make informed decisions and adjust the parameters to suit their specific needs. It also allows researchers with limited programming experience to perform the analysis with ease.

### Reanalysis of publicly available datasets

To demonstrate the capabilities of immunopipe, we have reanalyzed 9 publicly available datasets [10-18] (see Data availability). The datasets are provided with both scRNA-seq and scTCR-seq data. However, it is noteworthy that [15] contains an additional dataset with scRNA-seq data only (CD45+ cells), which can also be analyzed by immunopipe. The datasets cover not only cancer and cancer with therapy but also COVID-19 cases. The number of samples in each dataset ranges from 2 to 47 and individuals from 1 to 38; the number of cells varies from 6,438 to 194,519 and the number of matched TCR sequences from 5,642 to 77,030 (Table 1).

**Table 1.** Publicly available datasets reanalyzed by immunopipe.

| GEO | Condition | # individuals | # samples | # cells | # matched TCR sequences | Reference |
| --- | --- | --- | --- | --- | --- | --- |
| GSE144469 | Colitis and therapy | 22 | 22 | 75,569 | 68,760 | [15] |
| GSE176201 | COVID-19 lung tissue | 5 | 6 | 34,781 | 23,081 | [11] |
| GSE180268 | HPV and Head and Neck cancer | 6 | 19 | 53,303 | 26,844 | [12] |
| GSE114724 | Breast cancer | 3 | 5 | 28,341 | 24,039 | [10] |
| GSE161192 | LB dementia | 2 | 4 | 6,438 | 5,642 | [13] |
| GSE179994 | Anti-PD-1 therapy in lung cancer | 38 | 47 | 150,849 | 77,030 | [14] |
| GSE148190 | Skin cancer | 1 | 2 | 8,794 | 4,904 | [16] |
| GSE139555 | Anti-PD1 therapy | 14 | 32 | 194,519 | 67,700 | [17] |
| GSE145370 | Esophageal cancer | 7 | 14 | 108,226 | 35,449 | [18] |

The primary aim of the reanalysis is to demonstrate the capabilities of immunopipe, provide a reference for researchers to adjust the parameters, visualize the corresponding results, and facilitate the interpretation of the results. The reanalysis is not a reproduction of the analysis in the original publications, and the results may not be directly comparable. This is because the pre-processing steps are not fully revealed in some of the publications, and the analysis is not performed with the same parameters. Moreover, the tools used in the original analysis may have been updated since the publication, which can also lead to differences in the results (see also Discussion). The reanalysis is performed with the configuration files fed to immunopipe. Some studies use data other than scRNA-seq and scTCR-seq for their discoveries. In these cases, a minimal configuration file is provided to perform the analysis with only the necessary analyses enabled. If a study focuses on scRNA-seq and scTCR-seq analysis, the pipeline is configured to attempt to regenerate the figures in the publications. Configuration files, results, and reports are available in each repository (see Data availability).

## Discussion

Immunopipe uses data processed by 10X Genomics (https://www.10xgenomics.com/) CellRanger [48] by default. It can also take 10X-compatible scRNA-seq data, and scTCR-seq data from other platforms that can be loaded by immunarch [35]. In addition to the raw count and annotated contig files, a file with a serialized object, such as a Seurat object in RDS (https://rdrr.io/r/base/readRDS.html), h5Seurat (https://mojaveazure.github.io/seurat-disk/articles/h5Seurat-load.html) or AnnData [49], can be used as the input of immunopipe to adopt data from other platforms, even though it may take extra effort to load the data into Seurat objects. Moreover, it is beneficial to take the Seurat object as input for reproducibility. The reproducibility issue can be caused by inconsistent process steps and their parameters, and package versions. Regardless of the efforts, the reproducibility is still not guaranteed. For example, the irlba package [50] used in Seurat for dimension reduction is not deterministic, due to the numerical precision for manipulation on sparse matrix (discussed at https://github.com/bwlewis/irlba/issues/61). Incorporating packages such as rCASC [51], designed specifically for reproducibility, could be an option to consider; while adopting a Seurat object as input could be another enhancement for immunopipe in the future.

Currently, immunopipe only supports scTCR-seq data of αβ T cells. The γδ T cells are also a subset of T cells that express γδ TCRs instead of αβ TCRs [52, 53]. Even with less abundance (0.5–5% of all T-lymphocytes) than αβ T cells (65–70%), γδ T cells play a significant role in the immune system [54, 55]. Supporting γδ TCRs is an additional feature that can be added to immunopipe. The analytical methods employed for scTCR-seq data may be extrapolated for the analysis of single-cell B-cell receptor sequencing (scBCR-seq) data. The complexity of BCR data analysis is heightened by the occurrence of somatic hypermutation in B-cell receptors [56, 57], which can be addressed by distance-based clonotype analysis [58]. ScBCR-seq data analysis is supported by immunarch [35], which is used by immunopipe for scTCR-seq data analysis.

However, a B-cell selection process could be necessary to facilitate integrative analysis for scRNA-seq and scBCR-seq data.

In recent research studies, there has been an increase in the usage of additional downstream analysis based on scRNA-seq. This includes RNA velocity [59, 60], cell-cell communication [61], and gene regulatory networks [62]. These tools provide more detailed information on the biological processes happening at the single-cell level and can help researchers gain deeper insights into the mechanisms that underlie cellular behavior. The incorporation of these additional analysis modules into the immunopipe for scRNA-seq further empowers users to conduct comprehensive and insightful investigations, thereby facilitating a more comprehensive exploration of the data.

## Methods

Immunopipe is implemented on top of pipen (https://github.com/pwwang/pipen), a flexible and extensible pipeline framework written in Python (https://python.org). Its architecture facilitates the building of complex pipelines by breaking them down into smaller and reusable processes.

The extensibility of pipen enables a variety of plugins that add to its already comprehensive capabilities. Immunopipe benefits from two particularly essential plugins, including pipen-report (https://github.com/pwwang/pipen-report) which streamlines the process of organizing and generating reports for pipelines, and pipen-board (https://github.com/pwwang/pipen-board) which supplies a user-friendly web interface for configuring the pipeline, starting and monitoring the execution.

The implementation of each process is not restricted by programming languages, allowing for the incorporation of existing tools and packages. Immunopipe wraps both existing tools and in-house scripts to perform the analysis. Seurat [25] is the core tool for the pipeline, not only for the common practices of scRNA-seq data analysis but also for the integration of scRNA-seq and scTCR-seq data. The TCR clonal information is loaded into the Seurat object as metadata at the cell level and passed down to the downstream analysis. Enrichr [63] is used for enrichment analysis, and fgsea [64] is employed for GSEA. Automated cell type annotation is performed using scCATCH [65], sctype [66], celltypist [67], or hitype (https://github.com/pwwang/hitype), an enhanced version of sctype by introducing weights for markers. The scTCR-seq data is loaded into the pipeline by immunarch [35], which also performs basic analysis, including statistics on TCR clones, diversity metrics, gene usage, repertoire overlap, and clonotype tracking. Clustering by TCR sequences is conducted by GIANA [36] or clusTCR [37]. Metabolic landscape analysis is performed by the pipeline provided by [43], which is modified to fit into immunopipe. The physiochemical attribute analysis of the CDR3 amino acid sequences is performed using TiRP [47]. TESSA [23] is incorporated to identify phenotype-associated TCR repertoire. The rest of the analyses and visualizations are implemented by in-house scripts, including the T cell selection, visualization of gene expression patterns and other features, and exploration of cell distribution in different cell types, etc.

We provide the documentation for immunopipe online as a manual and a web interface to generate the configuration file, where the description of the options is shown immediately upon focus. To illustrate the utilization of immunopipe in the analysis of scRNA-seq and scTCR-seq data, we provide an illustrative example employing a publicly available dataset [17] (see Data availability). The example includes a minimal configuration file with only the necessary configuration for running the pipeline, which is ideal for a quick start, and other configuration files with different options enabled, which can be used as a reference for users to adjust the parameters to suit their specific needs. It also contains the results and report generated by the pipeline, which can be viewed online (see Data availability). We have also compiled a gallery of repositories containing the configuration files for the pipeline and the results of the reanalysis of 9 publicly available datasets (see Data availability). The gallery can be a useful resource and reference for the users with their analysis requirements.

## Data availability

The details about the example can be found at https://github.com/pwwang/immunopipe-example. The data used in the example is hosted at GEO (GSE139555). The report for the example generated by the minimal configuration file can be viewed at http://imp.pwwang.com/minimal/REPORTS/ and the report by the configuration file with all processes enabled is available at https://imp.pwwang.com/output/REPROTS/. The gallery of repositories for the reanalysis of the publicly available datasets can be found athttps://pwwang.github.io/immunopipe/gallery/.

## Code availability

The most recent source code for immunopipe is publicly available on GitHub (https://github.com/pwwang/immunopipe) and the documentation can be found at https://pwwang.github.io/immunopipe/. The docker image for immunopipe is also available on DockerHub (https://hub.docker.com/r/justold/immunopipe).

---

**Figure S1.**
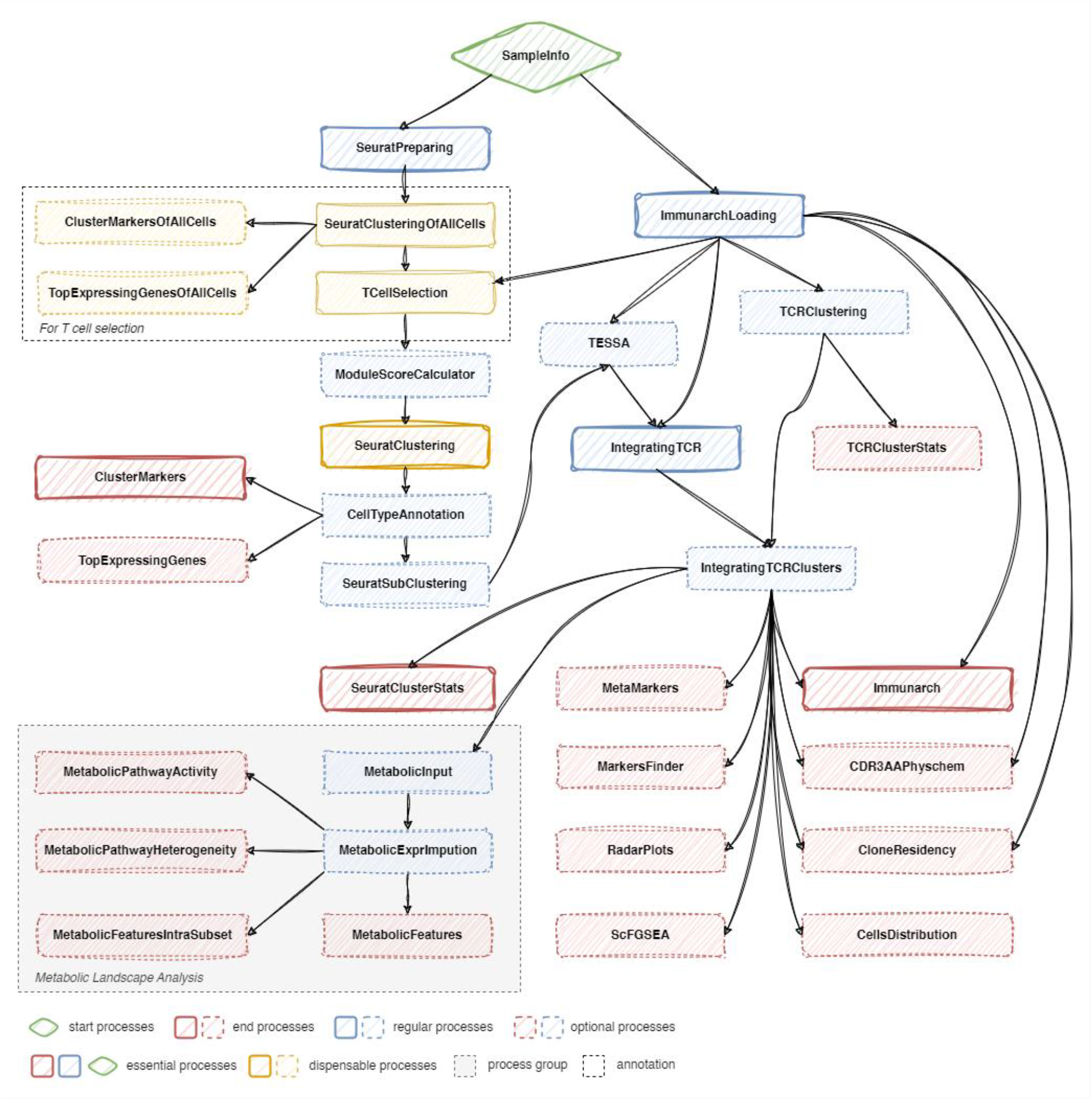
The flowchart of full analyses by immunopipe

**Figure S2.**
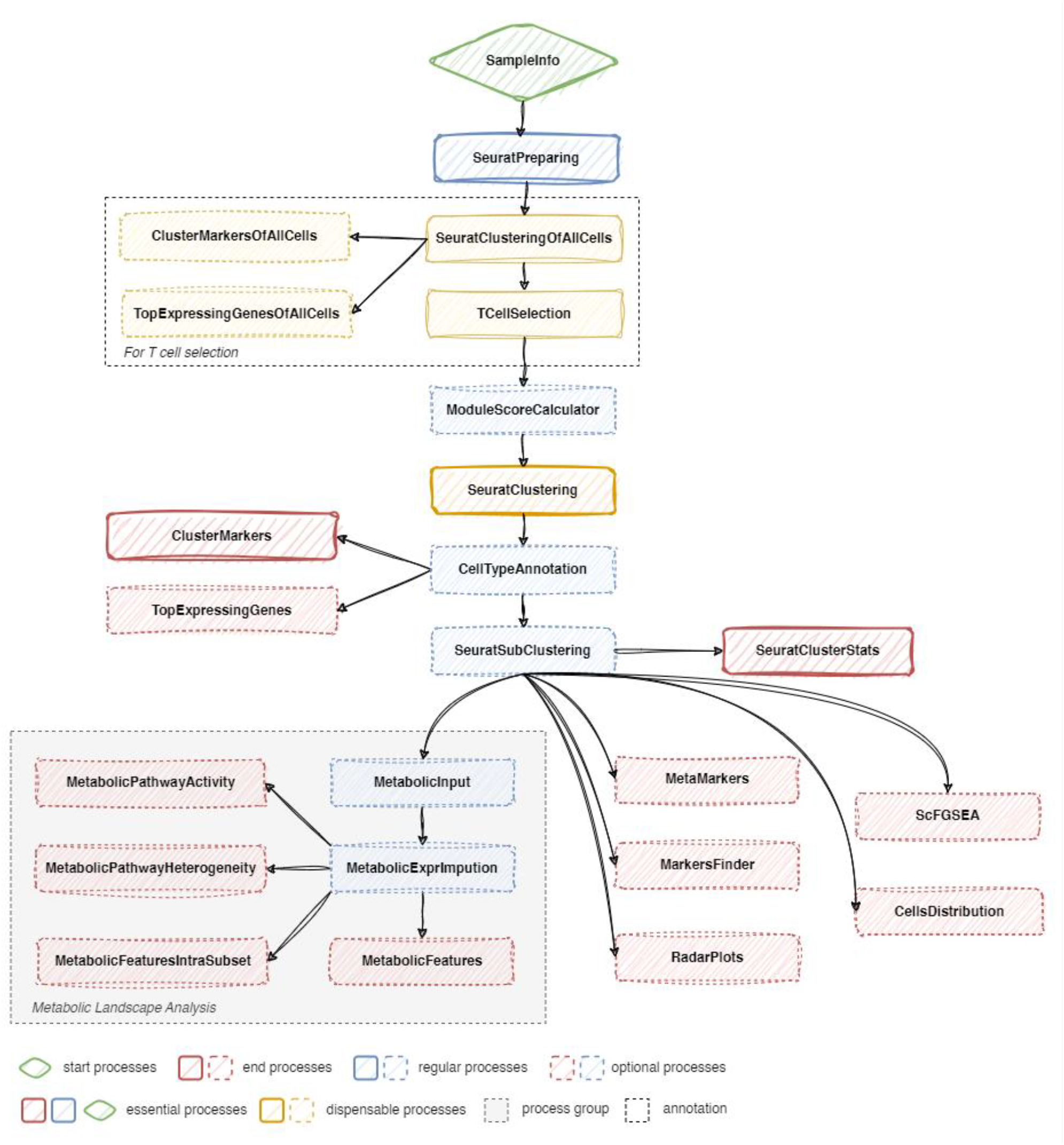
The flowchart of analyses with scRNA-seq data only by immunopipe

