## Supplementary figures and images for "Immunopipe: A comprehensive and flexible scRNA-seq and scTCR-seq data analysis pipeline"

### Figure S1

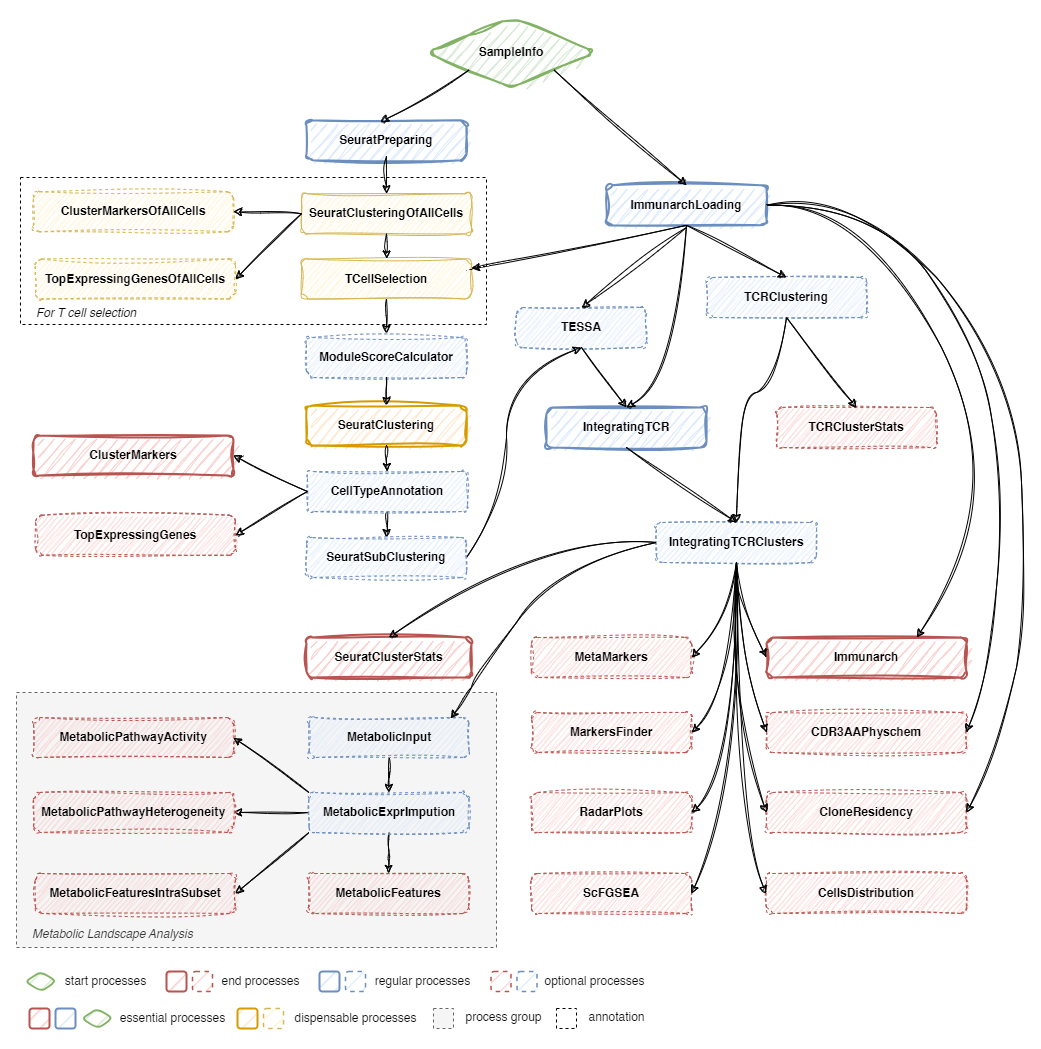

### Figure S2

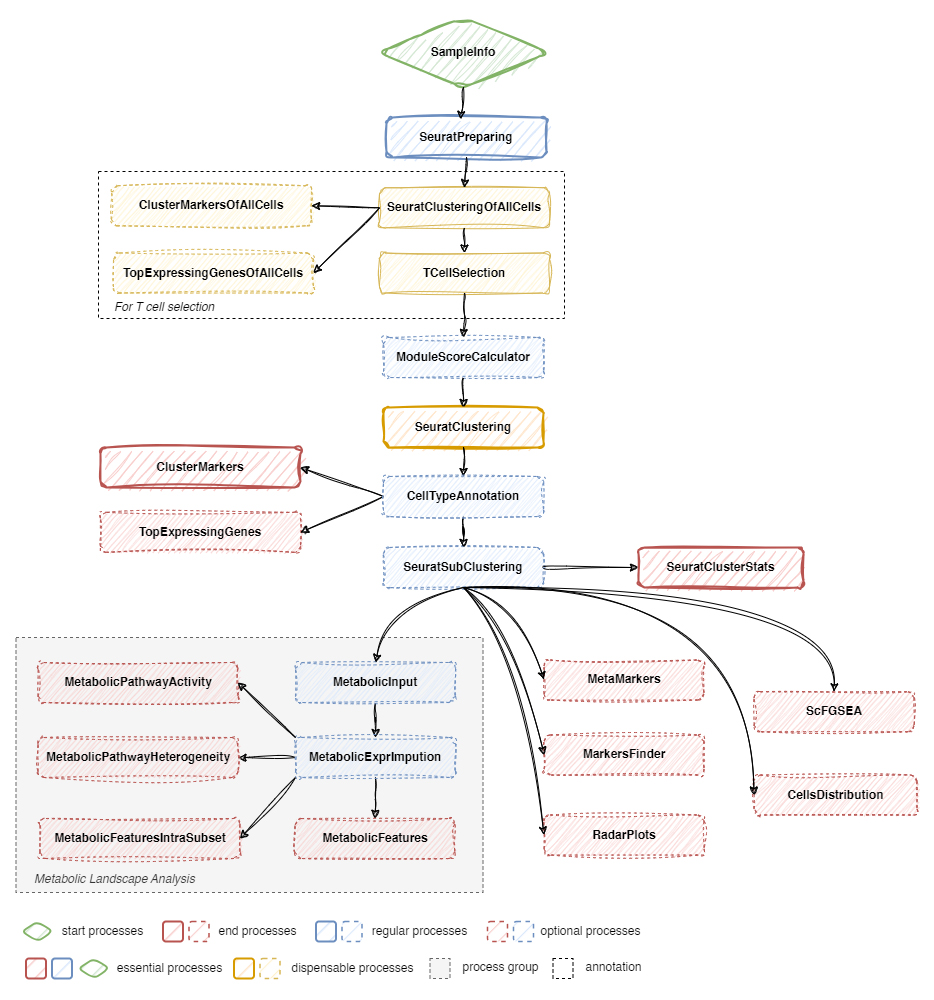
